# Mar1, an HMG-box protein, regulates *n*-alkane adsorption and cell morphology of the dimorphic yeast *Yarrowia lipolytica*

**DOI:** 10.1101/2024.03.22.586302

**Authors:** Chiaki Kimura-Ishimaru, Simiao Liang, Katsuro Matsuse, Ryo Iwama, Kenta Sato, Natsuhito Watanabe, Satoshi Tezaki, Hiroyuki Horiuchi, Ryouichi Fukuda

## Abstract

The dimorphic yeast *Yarrowia lipolytica* possesses an excellent ability to utilize *n*-alkane as a sole carbon and energy source. Although there are detailed studies on the enzymes that catalyze the reactions in the metabolic processes of *n*-alkane in *Y. lipolytica*, the molecular mechanism underlying the incorporation of *n*-alkane into the cells remains to be elucidated. Because *Y. lipolytica* adsorbs *n*-alkane, we postulated that *Y. lipolytica* incorporates *n*-alkane through direct interaction with it. We isolated and characterized mutants defective in adsorption to *n*-hexadecane. One of the mutants harbored a nonsense mutation in *MAR1* (<u>M</u>orphology and *n*-alkane <u>A</u>dsorption <u>R</u>egulator) encoding a protein containing a high mobility group box. The deletion mutant of *MAR1* exhibited defects in adsorption to *n*-hexadecane and filamentous growth on solid media, whereas the strain that overexpressed *MAR1* exhibited hyperfilamentous growth. Fluorescence microscopic observations suggested that Mar1 localizes in the nucleus. RNA-seq analysis revealed the alteration of the transcript levels of several genes, including those encoding transcription factors and cell surface proteins, by the deletion of *MAR1*. These findings suggest that *MAR1* is involved in the transcriptional regulation of the genes required for *n*-alkane adsorption and cell morphology transition.

**IMPORTANCE:** *Y. lipolytica*, a dimorphic yeast capable of assimilating *n*-alkane as a carbon and energy source, has been extensively studied as a promising host for bioconversion of *n*-alkane into useful chemicals and bioremediation of soil and water contaminated by petroleum. While the metabolic pathway of *n*-alkane in this yeast and the enzymes involved in this pathway have been well-characterized, the molecular mechanism to incorporate *n*-alkane into the cells is yet to be fully understood. Due to the ability of *Y. lipolytica* to adsorbs to *n*-alkane, it has been hypothesized that *Y. lipolytica* incorporates *n*-alkane through direct interaction with it. In this study, we identified a gene, *MAR1*, which plays a crucial role in the transcriptional regulation of the genes necessary for the adsorption to *n*-alkane and the transition of the cell morphology in *Y. lipolytica*. Our findings provide valuable insights that could lead to advanced applications of *Y. lipolytica* in *n*-alkane bioconversion and bioremediation.

## INTRODUCTION

The dimorphic yeast *Yarrowia lipolytica* possesses the abilities to utilize hydrophobic substrates, including *n*-alkane and triacylglycerol, as sole carbon and energy sources (1–3). Because of these characteristic features, *Y. lipolytica* has been investigated as a model organism to elucidate the metabolism of hydrophobic substrates and its regulation in yeasts (4–9). Furthermore, *Y. lipolytica* has attracted tremendous attention as a host for the bioconversion of hydrophobic substrates into various useful chemicals or for the bioremediation of soil and water contaminated by petroleum or oil (4, 10, 11).

In *Y. lipolytica*, incorporated *n*-alkane is hydroxylated to fatty alcohol by cytochrome P450 of the CYP52 family in the endoplasmic reticulum (ER). *Y. lipolytica* has 12 genes, *ALK1*–*ALK12*, encoding the CYP52-family P450s. Among these genes, *ALK1* and *ALK2* play primary roles in *n*-alkane oxidation, and the double deletion mutant of these genes exhibited a severe growth defect on media containing *n*-alkanes of various lengths as carbon and energy sources (12–16). Fatty alcohol is then oxidized to fatty aldehyde by fatty alcohol dehydrogenases or a fatty alcohol oxidase. *Y. lipolytica* has eight genes, *ADH1*–*ADH7* and *FADH*, encoding alcohol dehydrogenases and a gene, *FAO1*, encoding a fatty alcohol oxidase. The triple deletion mutant of *ADH1*, *ADH3*, and *FAO1* exhibited growth defects on medium containing fatty alcohol as a carbon and energy source, suggesting the essential roles of these genes in the oxidation of fatty alcohol (17, 18). Fatty aldehyde is further oxidized to fatty acid by fatty aldehyde dehydrogenases in the ER or peroxisome. Four fatty aldehyde dehydrogenase genes, *HFD1*– *HFD4*, are encoded in the genome of *Y. lipolytica*, which play critical roles in the oxidation of fatty aldehyde generated during the metabolism of *n*-alkane (19). Fatty acid is activated to acyl-CoA by acyl-CoA synthetases (20–24) and used for the synthesis of membrane and storage lipids or metabolized through β-oxidation in the peroxisome.

The transcription of a subset of genes involved in *n*-alkane assimilation, including *ALK1*, is activated in the presence of *n*-alkane and repressed in the presence of glycerol (12, 13, 25). The transcription of *ALK1* is regulated by the transcription activator complex composed of two basic helix-loop-helix transcription factors, Yas1 and Yas2, and by the Opi1-family transcription repressor Yas3 (14, 26–30).

Although the metabolic pathway of *n*-alkane and its regulation in *Y. lipolytica* have been well established, there is no clear understanding on the molecular mechanism by which *n*-alkane, a highly hydrophobic compound, is incorporated. Two mechanisms for facilitating the uptake of *n*-alkane have been proposed in *Y. lipolytica*. One is the utilization of biosurfactants/bioemulsifiers to solubilize hydrophobic substrates, and it has been reported that *Y. lipolytica* secretes biosurfactants/bioemulsifiers (31–33). The other mechanism is the uptake of *n*-alkane through direct interaction between the cell surface and *n*-alkane. *Y. lipolytica* adsorbs the droplet of *n*-alkane when cultured in liquid medium containing *n*-alkane, and it is possible that *Y. lipolytica* incorporates *n*-alkane through this interaction. Nevertheless, the molecular mechanism by which *Y. lipolytica* adsorbs the *n*-alkane droplet remains unclear.

*Y. lipolytica* is a dimorphic yeast that changes its cell morphology between the yeast form and pseudohyphal or hyphal form in response to various environmental conditions, including pH and nutrition (34). The cell morphology is regulated by transcription factors, such as Mhy1 and Hoy1, that promote the transition from the yeast to hyphal form and by transcription factors, such as Znc1, Tec1, Fts1, and Nrg1, that promote the transition from the hyphal to yeast form (35–40). Nonetheless, the entire picture of the transcriptional regulatory network of cell morphogenesis in *Y. lipolytica* remains to be established.

In this study, we isolated and characterized mutants that exhibited defects in *n*-alkane adsorption and identified a gene involved in the regulation of *n*-alkane adsorption and the morphological transition from the yeast to hyphal form in *Y. lipolytica*.

## RESULTS

### Isolation of mutants defective in *n*-alkane adsorption

To identify the genes involved in the adsorption of *Y. lipolytica* to *n*-alkane, we screened for mutants defective in *n*-alkane adsorption using the methods described in Experimental procedures. Briefly, the wild-type strain cultured in minimal medium containing glucose (SD) was collected and suspended in YNB medium containing *n*-hexadecane. After vigorous shaking, the suspension was centrifuged, and the pellet was collected to obtain cells that did not adsorb *n*-hexadecane. Then, a portion of the precipitated cells was inoculated into the SD medium and cultured again. This cycle was repeated five times to enrich the cells that lost the ability to adsorb to *n*-hexadecane by spontaneous mutations.

*Y. lipolytica* is a dimorphic yeast, and the wild-type *Y. lipolytica* strain forms rough and fluffy colonies on the SD plate because of the filamentous growth (Fig. 1A and B). However, when cells obtained by the enrichment culture of mutant strains that lost the ability of *n*-alkane adsorption were plated on SD plates, smooth colonies formed at high frequencies. Therefore, we isolated 80 strains that formed smooth colonies on SD plates after the enrichment culture. In this study, we characterized one of the obtained strains, Mutant 62.

**FIG 1.**
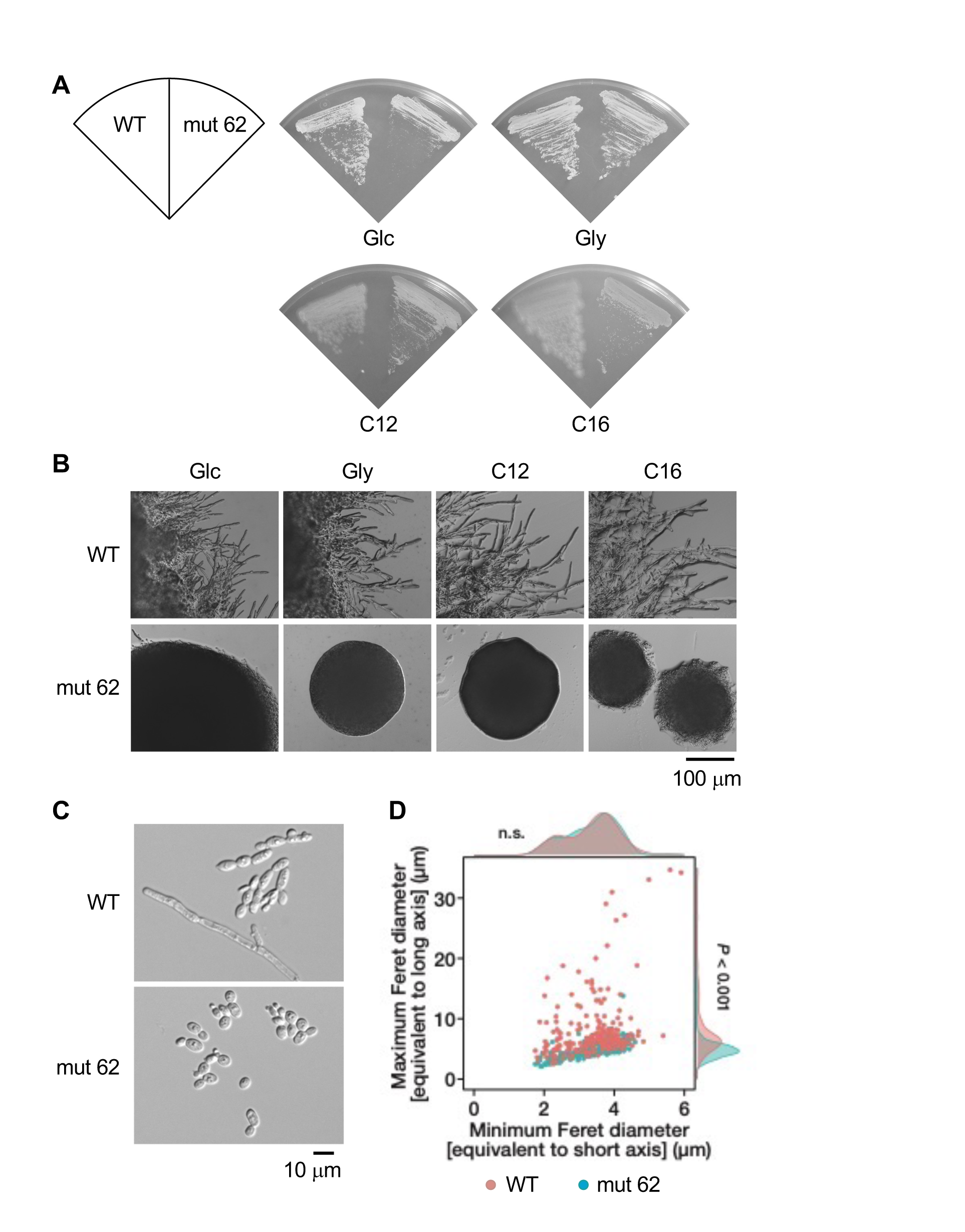
Growth and morphology of Mutant 62. (A) Growth of Mutant 62. The wild-type strain (WT) and Mutant 62 (mut 62) were cultured on the SD medium (Glc) for 3 days, SG medium (Gly) for 2 days, and on the YNB medium containing *n*-dodecane (C12) or *n*-hexadecane (C16) for 5 days. (B-D) Morphology of Mutant 62. (B) The wild-type strain and Mutant 62 were cultured as described in (A). The colony edges were observed under a microscope. Bar, 100 μm. (C,D) The wild-type strain and Mutant 62 were cultured in the liquid SD medium for 24 h. (C) The cell morphology of the wild-type strain and Mutant 62 was observed under a microscope. Bar, 10 μm. (D) The lengths of the long and short axes of the cells of the wild-type strain (red) and Mutant 62 (light blue) were measured. Scatter plots with marginal density plots of the long- and short-axes lengths of the cells are shown. The numbers of samples of the WT and mut 62 cells were 218 and 225, respectively.

We first analyzed the ability of Mutant 62 to adsorb *n*-hexadecane. The cells of the wild-type strain or Mutant 62 were suspended in YNB medium containing *n*-hexadecane. After vigorous shaking and centrifugation, the cells that adsorbed *n*-hexadecane were removed with the uppermost *n*-hexadecane layer and the supernatant. The optical densities of the suspension of pellet cells in the YNB medium were measured, and the ratios of pellet cells to the cells in the initial suspensions were determined. As depicted in Fig. 2, the ratio of the adsorption of Mutant 62 cells was significantly smaller than that of the wild-type cells, suggesting a defect in *n*-hexadecane adsorption.

**FIG 2.**
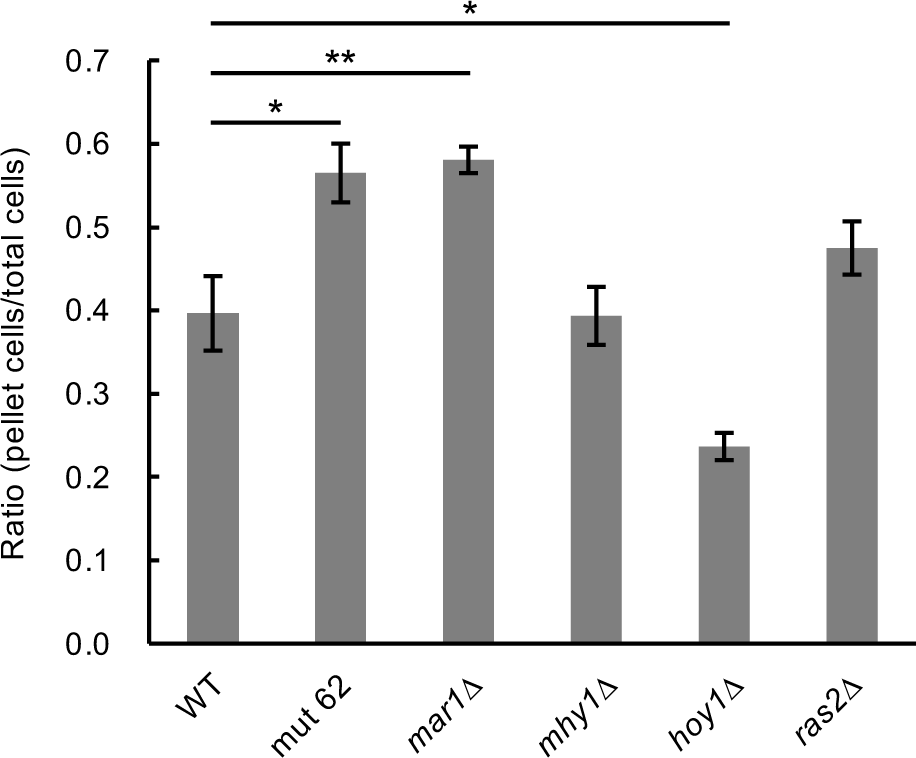
Adsorption of the *Y. lipolytica* strains to *n*-hexadecane. The adsorption of the wild-type strain (WT), Mutant 62 (mut 62), and the deletion mutants of *MAR1* (*mar1*Δ), *MHY1* (*mhy1*Δ), *HOY1* (*hoy1*Δ), and *RAS2* (*ras2*Δ) was analyzed as described in Experimental Procedures. The ratios of the cells that did not adsorb to *n*-hexadecane are shown. Bars indicate the mean of three biological replicates. Error bars represent S.E. Significant differences compared to the wild-type strain are indicated by asterisks (*, P < 0.05; **, P < 0.01; Dunnett’s test)

We next investigated the growth of Mutant 62 on solid media containing various carbon sources (Fig.1A and B). The wild-type strain exhibited filamentous growth around colonies on solid media containing glucose and glycerol and highly filamentous growth on media containing *n*-alkanes. Mutant 62 grew on medium containing glucose, glycerol, or *n*-alkanes but formed small colonies without filamentous growth on solid medium irrespective of the carbon source. In the liquid medium, the wild-type cells of *Y. lipolytica* exhibited yeast forms with various lengths of long axes and pseudohyphal forms (Fig. 1C). In contrast, the morphology of Mutant 62 cells was uniformly yeast form with short length of long axis (Fig. 1C). In fact, the long axes of Mutant 62 were smaller than those of the wild-type cells, whereas no significant differences were observed in the length of the short axes between the wild-type strain and Mutant 62 (Fig. 1D). Consistent with this, the areas occupied by the images of Mutant 62 cells in the micrographs were smaller than those of the wild-type cells, and the circularity of Mutant 62 cells was higher than that of the wild-type cells (Fig. S1). These findings suggest that Mutant 62 has defects in the regulation of cell morphology and filamentous growth, in addition to the adsorption to *n*-alkane.

To elucidate the relationship between cell morphology and *n*-alkane adsorption ability, we examined the *n*-alkane adsorption of the deletion mutants of *MHY1*, *HOY1*, and *RAS2*, which exhibit defects in the morphological transition from the yeast to hyphal form (35, 36, 41). The deletion mutant of *MHY1* exhibited a similar recovery level of cells in the pellet as that of the wild-type strain, and the deletion mutant of *HOY1* was less recovered in the pellet than that of the wild-type strain (Fig. 2). Therefore, no significant relationship was observed between cell morphology and *n*-alkane adsorption.

### Causative mutation for the defect in Mutant 62

Whole-genome sequencing of Mutant 62 was performed to determine the causative mutation(s) for the defect in *n*-alkane adsorption and the morphological transition of Mutant 62. We detected one single nucleotide substitution that altered the amino acid sequence of the translation product in the genome sequence of Mutant 62. The 265th nucleotide was substituted from C to T in the open reading frame (ORF) of *YALI0B02266g*, leading to the alteration of the 89th Gln codon to the termination codon. We termed *YALI0B02266g MAR1* (<u>M</u>orphology and *n*-alkane <u>A</u>dsorption <u>R</u>egulator), which was deduced to encode a 916-amino-acid protein of unknown function with a high mobility group (HMG) box (amino acid residues 152 - 219) (Fig. 3A), which has been proposed to be involved in DNA binding (42). The HMG box of *MAR1* product demonstrated significant similarities to those of the transcriptional repressors Rox1 of *Saccharomyces cerevisiae* (44% identity) and Rfg1 of *Candida albicans* (48% identity). Rox1 and Rfg1 are involved in the transcriptional repression of hypoxic genes (43) and the transcriptional regulation of filamentous growth (44, 45), respectively. The HMG box was deleted in the translation product of the *mal1* allele of Mutant 62.

**FIG 3.**
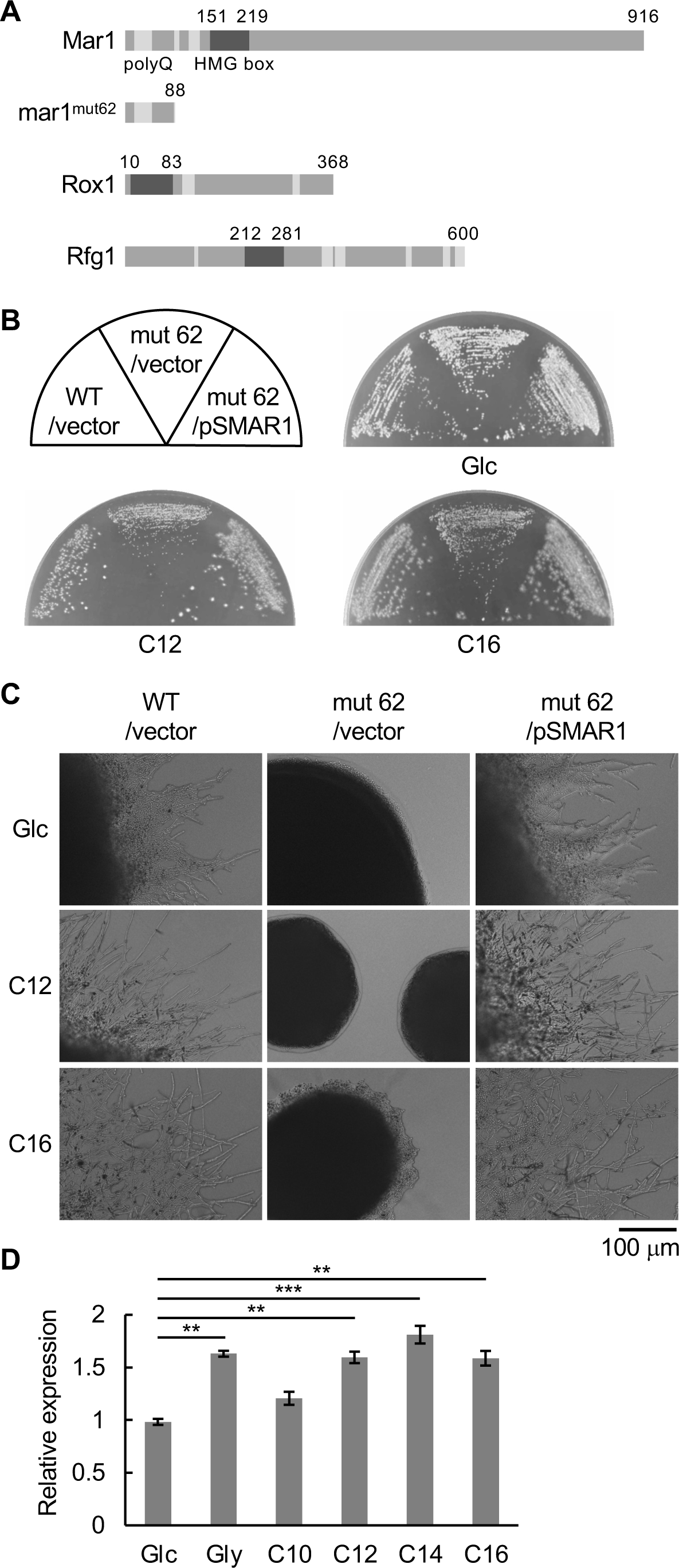
Structure of Mar1 and characterization of *MAR1*. (A) Schematic diagrams of Mar1 of the wild-type strain (Mar1) and Mutant 62 (mar1^mut62^), Rox1 of *S. cerevisiae*, and Rfg1 of *C. albicans*. The HMG boxes and Glutamine rich (polyQ) regions are indicated by dark and light grey, respectively. (B,C) Complementation of the defects of Mutant 62 by *MAR1*. (B) Growth of the wild-type strain (WT) and Mutant 62 (mut 62) harboring the empty vector or the plasmid to express *MAR1* (pSMAR1). Strains were cultured on the SD medium for 2 days and the YNB medium containing *n*-dodecane (C12) or *n*-hexadecane (C16) for 5 days. (C) Strains were cultured as described in (B). The colony edges were observed under a microscope. Bar, 100 μm. (D) qRT-PCR analysis of *MAR1*. The wild-type strain was cultured in the medium containing glucose (Glc), glycerol (Gly), *n*-decane (C10), *n*-dodecane (C12), *n*-tetradecane (C14), or *n*-hexadecane (C16) for 1 h. Cells were collected and the amount of *MAR1* mRNA was quantified as described in Experimental procedures. Bars indicate the mean of three biological replicates. Error bars represent S.E. Significant differences compared to the transcript level in the glucose-containing medium are indicated by asterisks (**, P<0.01; ***, P<0.001; Dunnett’s test).

To confirm that the mutation in *MAR1* is responsible for the phenotype of Mutant 62, a low-copy plasmid containing wild-type *MAR1* was introduced into Mutant 62, after which the growth was analyzed (Fig. 3B and C). The Mutant 62 expressing the wild-type *MAR1* exhibited filamentous growth on media containing glucose and *n*-alkanes, indicating that *MAR1* is the causative gene for the defect of the filamentous growth of Mutant 62.

The wild-type strain cultured in a glucose-containing medium for 24 h was shifted to a medium containing glucose, glycerol, *n*-decane, *n*-dodecane, *n*-tetradecane, or *n*-hexadecane as the carbon source and incubated for 1 h. The transcript levels of *MAR1*, analyzed by qRT-PCR, slightly increased after incubation in the medium containing glycerol, *n*-dodecane, *n*-tetradecane, or *n*-hexadecane compared with that with those in the medium containing glucose (Fig. 3D).

### Characterization of the deletion mutant of *MAR1*

Next, we constructed a deletion mutant of *MAR1* (*mar1*Δ). The *mar1*Δ strain exhibited a defect in *n*-hexadecane adsorption, similar to that of Mutant 62 (Fig. 2). *mar1*Δ was also defective in filamentous growth on solid media containing glucose or *n*-alkanes, similarly to Mutant 62 (Fig. 4A and B). *mar1*Δ exhibited a defect in pseudohyphal and hyphal growth in liquid media containing glucose or *n*-alkanes, and the long axes of *mar1*Δ cells were shorter than those of the wild-type strain cells, similar to that of Mutant 62 (Fig. 4C and D). Moreover, the areas occupied by the images of *mar1*Δ cells in the micrographs were smaller than those of the wild-type cells, and the circularity of *mar1*Δ cells was higher than that of the wild-type cells in these liquid media (Fig. S2). These findings suggest that *MAR1* is involved in the regulation of *n*-alkane adsorption and the morphological transition from yeast to pseudohyphal and hyphal forms. The deletion mutant of *MAR1* was slightly resistant to SDS on the rich medium containing glucose and the minimal medium containing *n*-hexadecane and slightly sensitive to Congo red on the medium containing *n*-hexadecane compared with the wild-type strain (Fig. S3), raising the possibility that *MAR1* is involved in the maintenance of the cell wall integrity.

**FIG 4.**
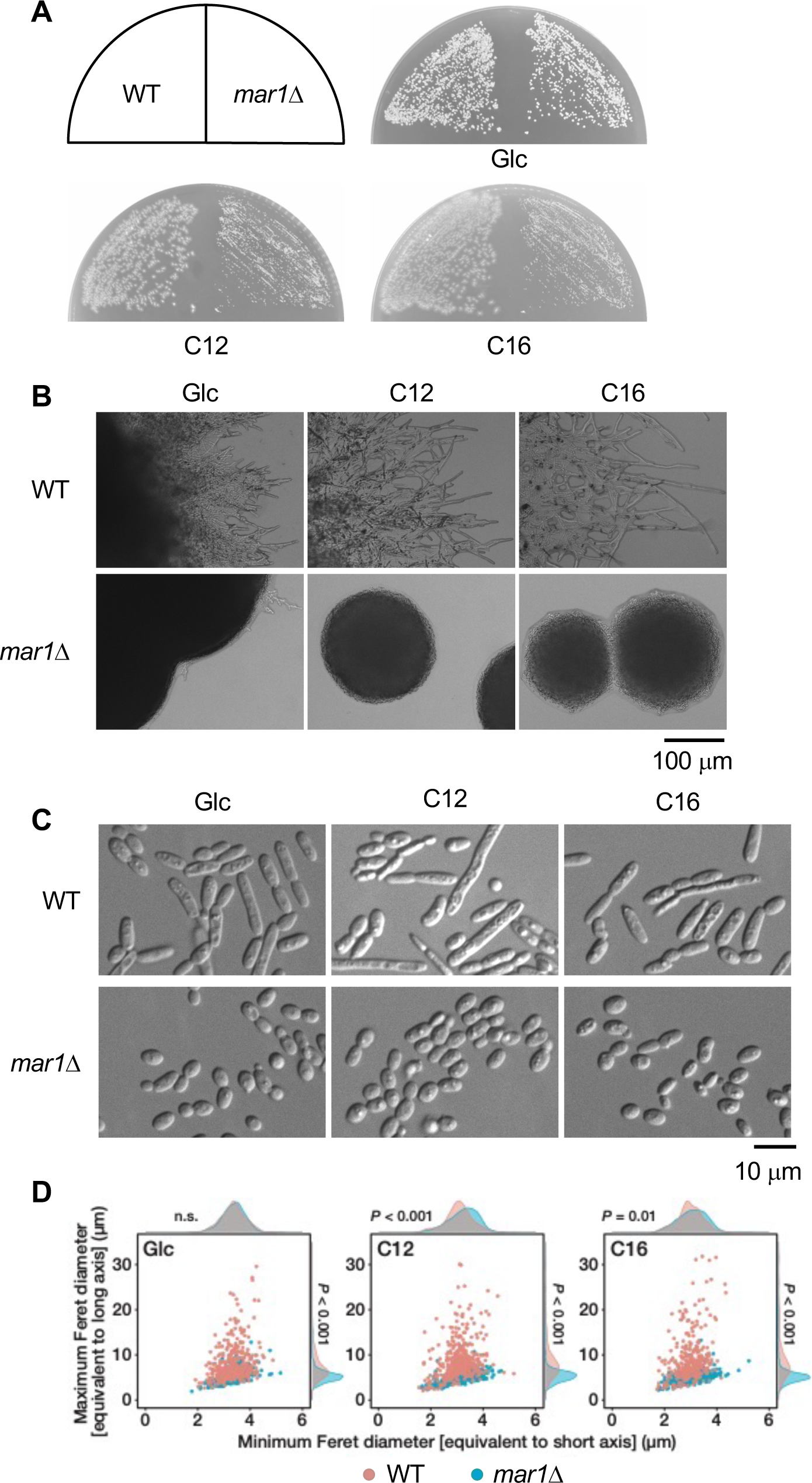
Growth and morphology of the deletion mutant of *MAR1*. (A) Growth of the deletion mutant of *MAR1*. The wild-type strain and *mar1*Δ were cultured on solid SD medium (Glc) for 3 days and on the YNB medium containing *n*-dodecane (C12) or *n*-hexadecane (C16) for 5 days. (B-D) Morphology of the deletion mutant of *MAR1*. (B) The wild-type strain and *mar1*Δ were cultured as described in (A). The colony edges were observed under a microscope. Bar, 100 μm. (C,D) The wild-type strain and *mar1*Δ were cultured in the liquid SD medium for 24 h and shifted to the SD medium (Glc) or the YNB medium containing *n*-dodecane (C12) or *n*-hexadecane (C16), after which they were further incubated for 6 h. (C) The cell morphology was observed under a microscope. Bar, 10 μm. (D) The lengths of the long and short axes of the cells of the wild-type strain (red) and *mar1*Δ (light blue) were measured. Scatter plots with marginal density plots of the long- and short-axes lengths of the cells are shown. The numbers of samples of the wild-type strain and mut 62 cells were 398 and 337 in glucose medium, 423 and 352 in C12 medium, and 315 and 370 in C16 medium, respectively.

### Subcellular localization of Mar1

The presence of an HMG box in Mar1 suggests that it binds to DNA, based on which we analyzed the localization of Mar1. We constructed a *Y. lipolytica* strain expressing Mar1 fused with EGFP at its N-terminus (EGFP-Mar1) under the *MAR1* promoter in its chromosomal location, as described in the Experimental Procedures section. This strain exhibited filamentous growth on the solid medium containing glucose, *n*-dodecane, or *n*-hexadecane (Fig. S4A and B), suggesting that EGFP-Mar1 has almost the same functionality as that of the wild-type Mar1 in *Y. lipolytica*. Western blot analysis of the extracts of cells cultured in the medium containing glucose or *n*-hexadecane using an anti-GFP antibody revealed a band with mobility similar to that expected from the molecular mass of the EFP-Mar1 fusion protein (126 kDa) (Fig. S4C). There was no significant difference in the amount of EGFP-Mar1 in the extracts of cells cultured in the medium containing glucose or *n*-hexadecane. We then observed the localization of EGFP-Mar1 by fluorescence microscopy in cells cultured to the logarithmic phase in the medium containing glucose and in those cultured in the medium containing *n*-hexadecane for an additional 3 h. The fluorescence signals of EGFP-Mar1 colocalized with those of Hoechst in cells cultured in both media (Fig. 5), suggesting that EGFP-Mar1 constitutively localizes in the nucleus.

**FIG 5.**
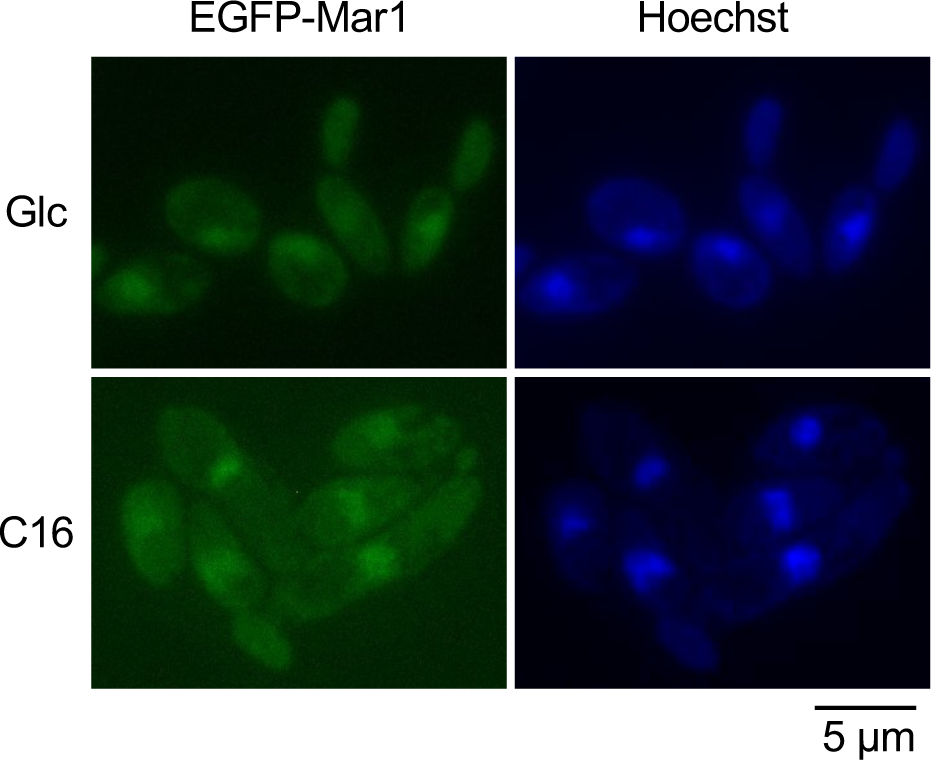
Subcellular localization of EGFP-Mar1. The wild-type strain expressing EGFP-Mar1 was cultured in the SD medium to logarithmic phase, after which it was shifted to the YNB medium containing *n*-hexadecane and incubated for 3 h. Localization of EGFP-Mar1 was observed as described in Experimental procedures. Bar, 5 μm.

### Genes under the regulation of Mar1

Because Mar1 contains the HMG box and localizes in the nucleus, it could function as a transcriptional regulator of the genes involved in *n*-alkane adsorption and morphological transition. To identify the genes whose transcript levels altered in the absence of Mar1, we conducted RNA-seq analysis of *mar1*Δ and its parental strain cultured in the medium containing glucose or *n*-hexadecane. We found that the transcript levels of 15 and 147 genes increased by more than 2-fold and those of 95 and 184 genes decreased by less than 1/2 in the deletion mutant of *MAR1* compared with those in the wild-type strain when cultured in media containing glucose and *n*-hexadecane, respectively (Fig. 6A; Table S1 and S2). The transcript levels of 6 genes increased in *mar1*Δ in both glucose-containing and *n*-hexadecane-containing media, whereas those of 25 genes decreased in both media (Table S3 and S4). Among the genes involved in the regulation of morphological transition in *Y. lipolytica*, the transcript levels of *RAS2*, *NRG1*, *FTS1*, *TUP1*, *SSN6*, *FTS2*, *RIM101*, *PHR1*, *BEM1*, *RAC1*, *BMH1*, *SIN3*, *ZNC1*, *TEC1*, *SCH9*, *RIM15*, and *ODC1* showed no significant changes by the deletion of *MAR1*, whereas the transcript level of *HOY1* decreased in *mar1*Δ cultured in the medium containing glucose or *n*-hexadecane. Moreover, the transcript level of *MHY1* decreased in *mar1*Δ cultured in the medium containing glucose. No significant differences were observed in the transcript levels of *MAR1* in the wild-type cells cultured in the medium containing glucose or *n*-hexadecane. Furthermore, the transcript levels of genes encoding various transcription factors and cell surface proteins showed alterations by the deletion of *MAR1*.

**FIG 6.**
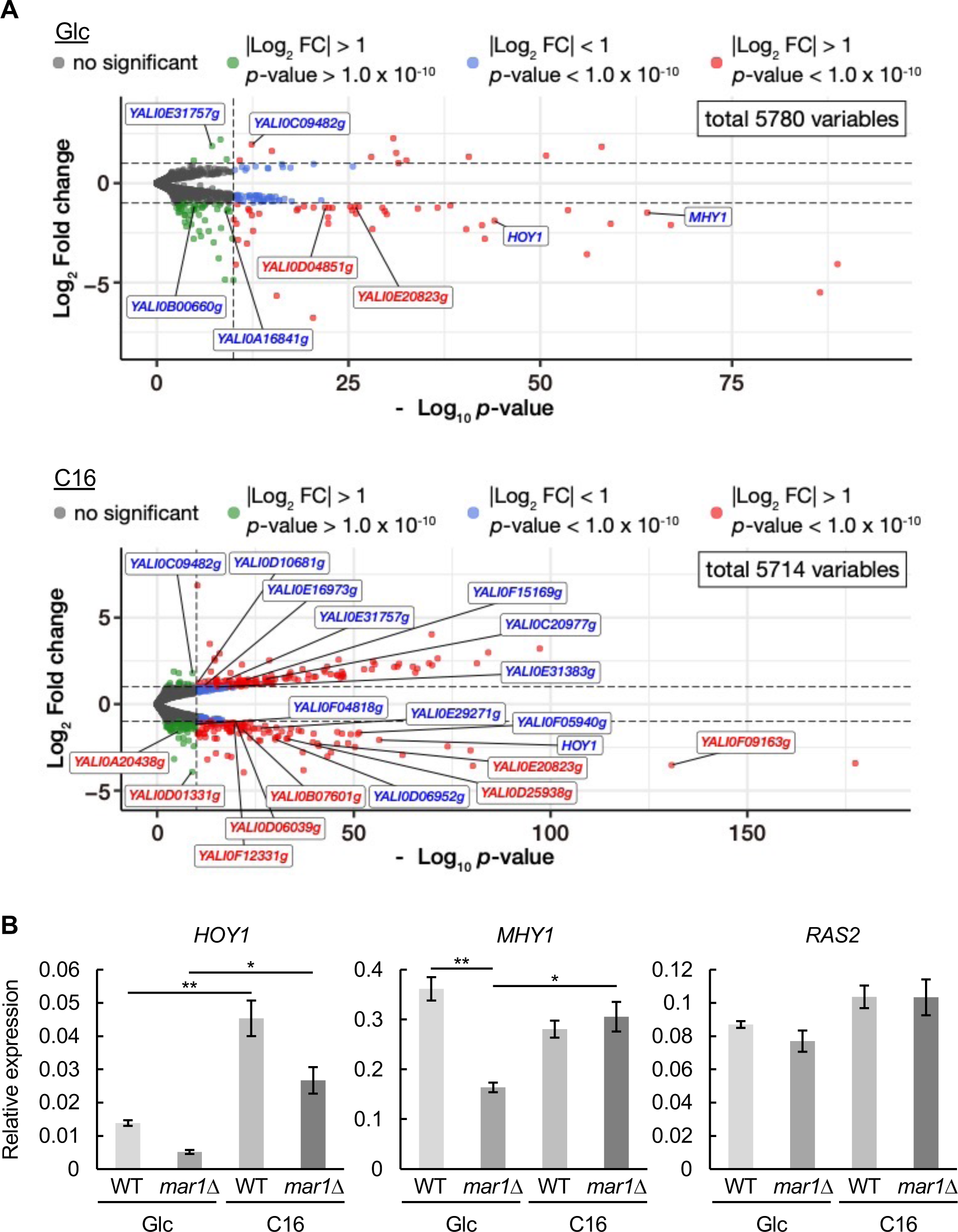
Transcriptional profile of the deletion mutant of *MAR1*. (A) RNA-seq analysis of *mar1*Δ cultured in the SD medium (Glc) or the YNB medium containing *n*-hexadecane (C16). Differential gene expression results are illustrated in the volcano plots, comparing *mar1*Δ to the wild-type strain. Each point represents a single gene. X-axis; −Log10 *p*-value, Y-axis; Log_2_ fold change in gene expression between *mar1*Δ and the wild-type strain. Positive values indicate upregulation in *mar1*Δ, while negative values indicate downregulation in *mar1*Δ. *Red* points; significantly upregulated/downregulated genes, *blue* points; significantly but slightly upregulated/downregulated genes, *green* points; upregulated/downregulated genes with low *p*-values, *gray* points; non-significantly differentially expressed genes, *horizontal dashed* lines; thresholds for log_2_ fold change (±1), *vertical dashed* line; threshold for statistical significance (*p*-value = 1.0 x 10^-10^). (B) qRT-PCR analysis of *HOY1*, *MHY1*, and *RAS2*. The wild-type strain and *mar1*Δ were cultured for 24 h at 30°C in the SD medium, and shifted to the SD medium (Glc) or the YNB medium containing *n*-hexadecane (C16), after which they were further incubated for 1 h. Cells were collected and the amounts of *HOY1*, *MHY1*, and *RAS2* mRNA were quantified according to Experimental procedures. Bars indicate the mean of three biological replicates. Error bars represent S.E. Statistically significant differences among transcripts are indicated by asterisks (*, P < 0.05; **, P < 0.01; Tukey-Kramer post-hoc test).

We analyzed the transcript levels of *HOY1* and those of a subset of genes involved in the regulation of morphological transition by qRT-PCR (Fig. 6B). Consistent with the results of the RNA-seq analysis, the transcript level of *HOY1* was lower in *mar1*Δ cultured in the medium containing glucose or *n*-hexadecane than in the wild-type strain. The transcript level of *MHY1* was also significantly lower in *mar1*Δ than in the wild-type strain cultured in media containing glucose, but not in media containing *n*-hexadecane. However, there was no significant difference in the amount of *RAS2* transcript between *mar1*Δ and the wild-type strain. These data suggest that Mar1 functions upstream of Hoy1 in the regulation of morphological transition from the yeast to hyphal form.

### Overexpression of *MAR1*

To determine the effect of the overexpression of *MAR1*, we constructed a plasmid to express *MAR1* under the constitutive promoter of *TEF1* encoding the translational elongation factor EF-1α (46) and introduced it into the wild-type strain. The transcript level of *MAR1* increased in the strain containing this plasmid compared with that in the wild-type strain containing the empty vector (Fig. 7A), although the level of overexpression was not so high. The growth of the strain harboring the *MAR1*-overexpressing plasmid retarded on the solid medium containing glucose or *n*-alkane compared with that of the strain harboring the vector (Fig. 7B). When the strain harboring the *MAR1*-overexpressing plasmid was cultured in the liquid medium containing glucose, filamentous growth was highly induced and the strain grew as mycelial pellets with its hyphae intertwined for at least 24 h (Fig. 7C and D), indicating that Mar1 promotes filamentous growth in *Y. lipolytica*.

**FIG 7.**
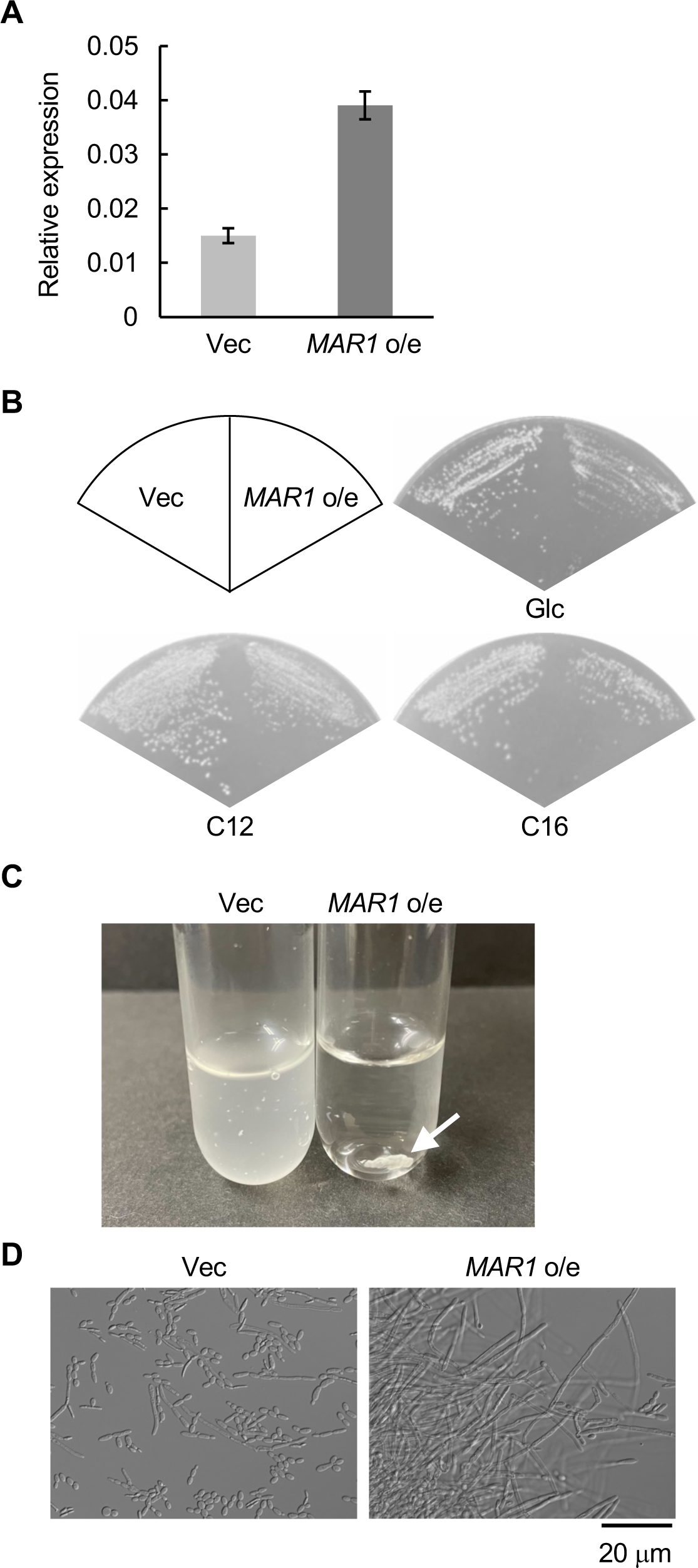
Overexpression of *MAR1*. (A) qRT-PCR analysis of *MAR1*. The wild-type strains containing the empty vector (Vec) and the *MAR1*-overexpressing plasmid (*MAR1* o/s) were cultured in the SD medium to logarithmic phase. Cells were collected and the amount of *MAR1* mRNA was quantified according to Experimental procedures. Bars indicate the mean of three biological replicates. Error bars represent S.E. (B) Growth of the strain containing the *MAR1*-overexpressing plasmid. The wild-type strains containing the empty vector and the *MAR1*-overexpressing plasmid were cultured on the SD medium for 3 days and the YNB medium containing *n*-dodecane (C12) or *n*-hexadecane (C16) for 5 days. (C,D) The wild-type strains containing the empty vector and the *MAR1*-overexpressing plasmid were cultured in the liquid SD medium for 24 h. (C) Cell cultures of the wild-type strains containing the empty vector and the *MAR1*-overexpressing plasmid. Arrow indicates the mycelial pellet of the *MAR1*-overexpressing strain. (D) Morphology of the *MAR1*-overexpressing strain were observed under a microscope. Bar, 20 μm.

## DISCUSSION

We identified *MAR1* encoding the HMG-box protein by characterizing a mutant defective in adsorption to *n*-hexadecane. The deletion mutant of *MAR1* exhibited defects in *n*-hexadecane adsorption and the morphological transition from the yeast to hyphal form, whereas the *MAR1*-overexpressing strain exhibited highly elongated filamentous growth. Moreover, the transcript levels of hundreds of genes altered in the deletion mutant of *MAR1*; in particular, the transcript level of *HOY1* decreased by the deletion of *MAR1*. These findings suggest that Mar1 is involved in the transcriptional regulation of genes involved in the adsorption of *n*-alkane and the morphological transition from the yeast to hyphal form.

The amino acid sequence of Mar1 demonstrated similarities to that of Rox1 of *S. cerevisiae* and that of Rfg1 of *C. albicans* in the HMG boxes. However, the size of Mar1 differs significantly from those of Rox1 and Rfg1 (Fig. 3A). Mar1 is a 916-amino-acid protein, whereas Rox1 and Rfg1 are proteins containing 368 and 600 amino acids, respectively. Moreover, Rox1 and Rfg1 have glutamine-rich (polyQ) regions similarly to Mar1, but their positions are different from that in Mar1. Mar1 has the polyQ regions at the N-terminal region of the HMG-box, whereas Rox1 and Rfg1 have the polyQ regions at the C-terminal regions of HMG boxes. Rox1 is a transcriptional repressor of hypoxic genes in *S. cerevisiae*, and the HMG domain of Rox1 was shown to bind to hypoxic operator sequences *in vitro* (43, 47). It was also demonstrated that the deletion of *ROX1* caused a defect in the pseudohyphal growth of the Σ1278b strain of *S. cerevisiae* (44), suggesting that Rox1 upregulated the transcription of genes involved in the morphological transition from the yeast to pseudohyphal form in the cells of the Σ1278b background of *S. cerevisiae*. In *C. albicans*, the deletion of *RFG1* increased filamentous growth on solid media, suggesting that Rfg1 functions as a transcriptional repressor of the genes involved in filamentous growth, which is in marked contrast to Mar1 and Rox1 (44, 45). It was also reported that *RFG1* overexpression decreased pseudohyphal growth in the Σ1278b strain of *S. cerevisiae* (44). These findings suggest that Rfg1 functions as a negative regulator of filamentous growth in not only *C. albicans* but also *S. cerevisiae*. In contrast to Rox1 in *S. cerevisiae*, Rfg1 did not appear to be involved in the transcriptional regulation of the hypoxic response in *C. albicans* (44). Therefore, one of the functions of the HMG box proteins in dimorphic yeasts could be to regulate cell morphology, but their roles vary by species.

The RNA-seq analysis suggested that the transcription of 15 and 147 genes was upregulated by more than 2-fold, and that of 95 and 184 genes was downregulated by less than 1/2 by the deletion of *MAR1* when cultured in media containing glucose and *n*-hexadecane, respectively (Fig. 6A; Table S1 and S2), implying that Mar1 is involved in the transcriptional regulation of a large number of genes. Furthermore, the transcript levels of 6 and 25 genes increased and decreased, respectively, in *mar1*Δ in both glucose- and *n*-hexadecane-containing media (Table S3 and S4). Among these genes, the transcript level of *HOY1*, which is involved in morphological transition, significantly decreased by the deletion of *MAR1* (Fig. 6A and B). The transcript levels of *MAR1* (*YALI0B02266g*) and *HOY1* were repressed by Tup1 and Ssn6, both of which are transcriptional repressors involved in filamentous growth (40). These data suggest that Mar1 functions downstream of Tup1 and Ssn6 and upstream of Hoy1 in morphological regulation in *Y. lipolytica*. Interestingly, Rfg1 functions in Tup1-dependent and -independent pathways in *C. albicans* (48). The roles of the genes, whose transcript levels increased or decreased in both glucose- and *n*-hexadecane-containing media, other than *HOY1*, remain to be determined.

The transcript level of *HOY1* in *mar1*Δ cells cultured in the *n*-hexadecane-containing medium was higher than that in the wild-type cells cultured in the glucose-containing medium, although *mar1*Δ exhibited defects in hyphal growth on the solid medium containing glucose or *n*-alkanes, and their long axes were shorter than those of the wild-type strain cells in liquid media containing glucose and *n*-alkanes. Hence, it is plausible that there exist pathway(s) other than that mediated by Hoy1 to regulate cell morphology downstream of Mar1. Furthermore, because the deletion mutant of *HOY1* exhibited no defect in *n*-hexadecane adsorption (Fig. 2), Hoy1 is not involved in the regulation of *n*-alkane adsorption. Therefore, *n*-alkane adsorption regulated downstream of Mar1 should be a pathway that does not involve Hoy1. Interestingly, the transcript levels of several genes encoding cell surface proteins decreased by the deletion of *MAR1* (Fig. 6); these genes could be involved in determining the degree of cell surface hydrophobicity.

In *C. albicans*, the cell wall proteome profile changes upon hyphal induction (49). Cell surface hydrophobicity should be important for *n*-alkane adsorption, and it is possible that there exists an intimate relationship between cell morphology and hydrophobicity of the cell surface in yeasts. In fact, it has been demonstrated that cell surface hydrophobicity is altered by filamentation in *Candida tropicalis* (50). Several drugs have also been reported to reduce cell surface hydrophobicity and inhibit filamentous growth in *C. albicans* (51–60). Therefore, it is possible that there exist mechanisms that simultaneously regulate cell surface hydrophobicity and cell morphology in dimorphic yeasts and that Mar1 is involved in their regulation in *Y. lipolytica*.

The deletion mutant of *MAR1* could grow in the presence of *n*-alkanes, although it formed smaller colonies on the solid medium containing *n*-alkanes (Fig. 4A and B). This could be due to the defect of *mar1*Δ in filamentous growth; therefore, it is not clear whether the ability to adsorb *n*-alkane is essential for the utilization of *n*-alkane. The deletion mutant of *MAR1* still had a partial ability to adsorb *n*-hexadecane (Fig. 2), and it is possible that there exist other mechanisms to adsorb *n*-alkane than that regulated by *MAR1*.

The RNA-seq analysis revealed much higher transcript level of *TEF1* than that of *MAR1* (Fig. S5), suggesting that the promoter of *TEF1* is much stronger than that of *MAR1*. When *MAR1* was expressed using the promoter of *TEF1*, the transcript level of *MAR1* increased, but not to the extent anticipated from the transcript level of *TEF1*. *MAR1* overexpression exerts negative effects on the growth of *Y. lipolytica*, as illustrated in Fig. 7B, and the mechanism that controls the transcript level of *MAR1* could exist in *Y. lipolytica*.

We isolated several mutants defective in *n*-alkane adsorption, in addition to Mutant 62. These mutants could harbor mutations in genes involved in *n*-alkane adsorption under the control of Mar1 or in regulatory pathways independent of Mar1 regulation. An analysis of these mutants might provide a clearer picture of the mechanisms of *n*-alkane adsorption and metabolism by *Y. lipolytica*, as well as the relationship between *n*-alkane metabolism and cell morphology.

## MATERIALS AND METHODS

### Yeast strains and growth conditions

Yeast strains used are listed in Table 1. *Y. lipolytica* strain CXAU1 (*ura3 ade1*) derived from CX161-1B (ATCC32338, *ade1*) was used as a wild-type strain (12).

**TABLE 1.** Yeast strains used in this study.

| Strain | Genotype | Source of reference |
| --- | --- | --- |
| CXAU1 | *MATA ade1 ura3* | 12 |
| $\Delta mar1$ | CXAU1 *mar1* | This study |
| $\Delta mhy1$ | CXAU1 *mhy1::ADE1* | This study |
| $\Delta ras2$ | CXAU1 *ras2::ADE1* | This study |
| $\Delta hoy1$ | CXAU1 *hoy1* | This study |
| GMA | CXAU1 *mar1::EGFP-MAR1* | This study |

Construction of the deletion mutants of *MAR1* and *HOY1* and the GMA strain, which produces Mar1 tagged with EGFP at its N-terminus, were carried out with the pop-in/pop-out method (15). *RAS2* and *MHY1* were deleted using *ADE1* as a marker. Correct integrations of the deletion cassettes were confirmed by PCR.

Yeast strains were cultured in YPD medium [2% Hipolypepton (FUJIFILM), 1% Yeast extract (Becton, Dickinson and Company), and 2% glucose] or minimal medium described below. Minimal medium was prepared by adding an appropriate carbon source to YNB [0.17% yeast nitrogen base without amino acids and ammonium sulfate (Becton, Dickinson and Company), 0.5% ammonium sulfate] as follows: 2% (w/v) glucose (SD); 2% (w/v) glycerol (SG); 2% (v/v) *n*-alkanes. Uracil (24 mg/l) and/or adenine (24 mg/l) were/was added, if necessary. For solid media, 2% agar was added. *n*-Alkanes were supplied in the vapor phase to YNB solid media. A piece of filter paper was soaked with *n*-alkanes and placed on the lid of a Petri dish, which was sealed and kept upside down. Yeast cells were grown at 30℃.

### Plasmids

Plasmids used in this study are shown in Table 2, and sequences of used primers are listed in Table 3.

**TABLE 2.** Plasmids used in this study.

| Plasmid | Descriptions | Reference or source |
| --- | --- | --- |
| pBluescript II SK (+) | A cloning vector | Agilent |
| pSUT5 | A low copy YCp plasmid carrying *URA3* | 26 |
| pSAT4 | A low copy YCp plasmid carrying *ADE1* | 12 |
| pBURA3 | pBluescript II SK (+) carrying *URA3* | 15 |
| pBADE1-RAS2PT | pBluescript II SK (+) carrying *RAS2* deletion cassette | This study |
| pBADE1-MHY1PT | pBluescript II SK (+) carrying *MHY1* deletion cassette | This study |
| pSMAR1 | pSUT5 carrying *MAR1* | This study |
| pBUMAR1 | A plasmid for deletion of *MAR1* by pop-in-pop-out method | This study |
| pBS-EGFP | pBluescript II SK (+) carrying EGFP | 19 |
| pBUGFP-MAR1 | A plasmid for construction of GMA | This study |
| pBUHOY1 | A plasmid for deletion of *HOY1* by pop-in-pop-out method | This study |
| pSU-TEF1p-MAR1 | A plasmid to overexpress *MAR1* under *TEF1* promoter | This study |

The plasmid to express *MAR1* was constructed as follows: the ORF of *MAR1* with the 5’- and 3’-adjacent regions were amplified from CXAU1 total DNA by PCR using primers MAR1P-F and MAR1T-R. Obtained fragment was digested with BamHI and EcoRI, and cloned into pSUT5 digested with same restriction enzymes, resulting in pSMAR1.

The deletion cassettes for *MAR1* and *HOY1* were constructed as follows. The 5’- and 3’-adjacent regions of these genes were amplified by PCR using primers, MAR1P-F, MAR1P-R, MAR1T-F, and MAR1T-R for *MAR1* and HOY1P-F, HOY1P-R, HOY1T-F, and HOY1T-R for *HOY1* and cloned into BamHI-EcoRI sites of pBURA3, to obtain pBUMAR1 and pBUHOY1, respectively. The deletion cassettes for these genes were obtained by digestion of pBUMAR1 with Van91I and pBUHOY1 with BspT140I.

The deletion cassettes for *RAS2* and *MHY1* were constructed as follows. The 5’- and 3’-untranslated regions (UTRs) of *RAS2* was amplified with primers RAS2P-F, RAS2P-R, RAS2T-F, and RAS2T-R, and digested with ApaI and BamHI, and BamHI and XbaI, respectively. These fragments were cloned into ApaI-XbaI sites of pBluescript II SK(+), resulting in pBRAS2PT. The 5’- and 3’-UTRs of *MHY1* was amplified with primers MHY1P-F, MHY1P-R, MHY1T-F, and MHY1T-R, and digested with EcoRI and BamHI, and BamHI and XbaI, respectively. These fragments were cloned into EcoRI-XbaI sites of pBluescript II SK(+), resulting in pBMHY1PT. The fragment containing *ADE1* was obtained by the digestion of pSAT4 (12) with BamHI and inserted into BamHI sites of pBRAS2PT and pBMHY1PT, yielding the plasmids pBADE1-RAS2PT and pBADE1-MHY1PT, respectively. The deletion cassettes were obtained by digestion of pBADE1-RAS2PT with ApaI and XbaI and pBADE1-MHY1PT with EcoRI and XbaI.

The plasmid pBUGFP-MAR1 carrying a cassette to express Mar1 fused with EGFP at its N-terminus in its chromosomal location was constructed as follows: the 5’-UTR and 5’-portion of the ORF of *MAR1* was amplified with primers MAR1P-F and MAR1-1000-R. The amplified fragment was digested with BamHI and KpnI and cloned into BamHI-KpnI sites of pBURA3. This plasmid was linearized by PCR using primers MAR1-pro and MAR1-ORF. The fragment carrying EGFP ORF was amplified by PCR from the plasmid pBS-GFP (19) using the primers EGFP-MAR1-F and EGFP-MAR1-R. These fragments were ligated with seamless ligation cloning extract (SLiCE) method (61), yielding the plasmid pBUGFP-MAR1. The cassette to express Mar1 fused with EGFP was obtained by digestion of pBUGFP-MAR1 with MluI.

The plasmid to overexpress *MAR1* was constructed as follows: the ORF of *MAR1* was amplified from CXAU1 total DNA by PCR using primers F-EcoRI-MAR1 and R-KpnI-MAR1. Obtained fragment were digested with EcoRI and KpnI, and cloned into pSUTEF1 (16) digested with same restriction enzymes, resulting in pSU-TEF1p-MAR1.

### Transformation of *Y. lipolytica*

*Yarrowia lipolytica* was transformed by electroporation as described previously (12)..

### Screening for mutants defective in adsorption to *n*-hexadecane

The CXAU1 strain precultured in the SD medium for 2 days was inoculated to 200 mL SD medium at an initial OD_600_ = 0.05 and cultured for 30 h, after which cells were collected by centrifugation. The precipitated cells were suspended in 20 mL YNB medium with 2 mL *n*-hexadecane. After vigorously shaking, the suspension was centrifugated, and the pellet was collected to obtain the cells that did not adsorb to *n*-hexadecane. The precipitated cells were inoculated to 200 mL SD medium at an initial OD_600_ = 0.05 and cultured for 30 h. After this cycle was repeated five times, the precipitated cells were plated on the SD medium and incubated for 3 days.

### Measurement of adsorption to *n*-hexadecane

Each strain was cultured in the SD medium to exponential growth phase, and the cells of approximately 15 OD_600_ units were collected by centrifugation. The precipitated cells were suspended in 10 mL YNB medium, and the initial OD_600_ was measured. After the addition of 1 mL *n*-hexadecane and vigorous shaking, the cell suspensions were centrifugated. The supernatants were removed, and the precipitated cells were suspended in 5 mL YNB media, after which the OD_600_ of these suspensions were measured. The *n*-alkane adsorption was calculated as follows: ratio (pellet cells/total input cells) = final OD_600_ of suspensions × 5 / initial OD_600_ of suspensions × 10.

### Microscopy

Microscopic images were acquired using BX52 microscope equipped with a microscope digital camera DP80 (Olympus, Tokyo, Japan). Long and short axes of cells were measured using ImageJ (Wayne Rasband), and the ratios of two axes were calculated. Nuclei were stained with Hoechst (FUJIFILM). For Hoechst staining, cells were collected, washed with PBS, and fixed in 70% ethanol for 15 min. Cells were washed twice and stained in 10 μg/ml Hoechst.

### Genome sequence analysis of mutant 62

Genome sequencing was performed by BGI Japan (Kobe, Japan) using Illumina HiSeq system with 150 base-paired end reads.

### RNA-seq analysis

The extracted total RNA samples were subjected to strand-specific mRNA-seq analysis using the DNBseq by BGI Japan (Kobe, Japan). mRNAs with poly-A tails were enriched using oligo dT beads and the enriched mRNAs were then fragmented into small pieces. The fragmented mRNAs were reverse-transcribed into first-strand cDNA using random primers. Subsequently, second-strand cDNA synthesis was performed using dUTP. The synthesized cDNA underwent end-repair and 3’-adenylation, followed by ligating adaptors to the 3’-adenylated ends of the cDNA fragments. The cDNAs were treated with uracil-DNA-glycosylase, and treated cDNA then underwent PCR amplification. After confirming the library’s quality, they were circularized and DNA nanoballs (DNBs) were generated in preparation for sequencing on the DNBseq platform.

### Bioinformatic process of RNA-seq data

Raw data with adapter sequences or low-quality sequences was filtered by SOAOnuke software (SOAPnuke software filter parameters: “ -n 0.01 -l 20 -q 0.4 --adaMR 0.25 --ada_trim --polyX 50 -- minReadLen 150”) (62). The obtained FASTQ files, representing the clean reads after the preprocessing steps, were registered in the DDBJ Sequence Read Archive (DRA) as the accession number: BioProject ID, PRJDB17495; BioSampleID, SAMD00736876 −SAMD00736879.

The FASTQ data were aligned to the reference genome of *Y. lipolytica* CLIB122 (assembly ASM252v1) using the Salmon software (63). All analyses were performed using R Statistical Software (version 4.3.1; R Core Team, 2023) and RStudio (v2023.06.1+524; Posit team, 2023), with R packages: tidyverse (v2.0.0; Wickham H, 2019) (64), DESeq2 (v1.40.2; Love MI et al., 2014) (65), and apeglm (v1.22.1; Zhu A et al., 2018) (66). Drawing volcano plots was performed using R packages: ggplot2 (v3.4.4; Wickham H, 2016) and EnhancedVolcano (v1.18.0) (67).

### Quantitative real time PCR (qRT-PCR)

Total RNAs were extracted using ISOGENII according to the manufacturer’s instructions. DNA was removed using DNaseI (Takara Bio Inc.). Reverse transcription of RNA was performed using PrimeScript^TM^ reagent Kit (Perfect Real Time) (Takara Bio Inc.) according to the manufacturer’s instructions. Real-time quantitative PCR was performed with primers, MAR1_U and MAR1_L *MAR1*, HOY1_U and HOY1_L for *HOY1*, MHY1_U and MHY1_L for *MHY1*, RAS2_U and RAS2_L for *RAS2*, and HHT1_U and HHT1_L for *HHT1* (Table 3) using SYBR^®^ Premix Ex Taq^TM^ (Perfect Real Time) (Takara Bio Inc.) according to the manufacturer’s instructions.

### Western blot analysis

Cells were collected and suspended in the breaking buffer [Tris-HCl (pH 7.5), 100 mM KCl, 10% (w/v) glycerol, 1 mM DTT, and 1% (v/v) Protease inhibitor cocktail for use with fungal and yeast extracts, DMSO solution (Sigma-Aldrich)] and disrupted with glass beads using Multi-Beads Shocker (YASUI KIKAI). After centrifugation at 1,000 x g for 10 min at 4°C twice, the supernatant was used as the whole cell extract. The whole cell extract was subjected to sodium dodecyl sulfate-polyacrylamide gel electrophoresis (SDS-PAGE) and proteins were transferred to a nitrocellulose membrane. EGFP-Mar1 was detected with Living Colors A.v. Monoclonal Antibody (JL-8) (Clontech). Anti-mouse IgG was used as a secondary antibody.

## ACKNOWLEDGMENTS

This work was partly supported by JSPS KAKENHI Grant (17K07710) and the Institute for Fermentation, Osaka (IFO): IFO research grant GK-2023-2-076. This work was performed using the facilities of the Agro-Biotechnology Research Center of The University of Tokyo.

Funding information

## AUTHOR CONTRIBUTION

The conception or design of the study: RI and RF; the acquisition, analysis, or interpretation of the data: CKI, SL, KM, RI, KS, NW, ST, HH, and RF; and writing of the manuscript: HH and RF.

## AVAILABILITY OF DATA

The data that support the findings of this study are available from the corresponding author upon reasonable request.

